# Machine Learning Approach to Predicting Stem-Cell Donor Availabilitys

**DOI:** 10.1101/242719

**Authors:** Adarsh Sivasankaran, Eric Williams, Mark Albrecht, Galen E. Switzer, Vladimir Cherkassky, Martin Maiers

## Abstract

The success of Unrelated Donor stem-cell transplants depends not only on finding genetically matched donors but also on donor availability. On average 50% of potential donors in the NMDP database are unavailable for a variety of reasons, after initially matching a patient, with significant variations in availability among subgroups (e.g., by race or age). Several studies have established univariate donor characteristics associated with availability. Individual consideration of each applicable characteristic is laborious. Extrapolating group averages to individual donor level tends to be highly inaccurate. In the current environment with enhanced donor data collection, we can make better estimates of individual donor availability. In this study, we propose a Machine Learning based approach to predict availability of every registered donor, to be used during donor selection and reduce the time taken to complete a transplant.

## 1. Introduction

Stem cell transplants are a curative therapy for a number of malignant and non-malignant hematological disorders. An ideal candidate donor for a stem cell transplant is an HLA (Human Leukocyte Antigen) matched sibling (Ballen et al. 2008). Unfortunately, only 30% of patients can find HLA compatible donors within their families (Besse et al. 2016). The remaining 70% rely on either Unrelated Donors (URDs), Cord Blood Units (CBUs) or Haploidentical Donors for a successful transplant. URD and CBU searches are performed with the assistance of public donor registries. The National Marrow Donor Program^®^ (NMDP) provides access to the US Be The Match Registry^®^ as well as several other domestic and international registries. The NMDP database has over 20 million adult donors, and over 450,000 cord blood units.

HLA compatibility for URD searches is determined by an HLA matching algorithm, such as HapLogic™ (Dehn et al. 2016) used by the NMDP. HLA compatible donors are then displayed to physicians (and/or clinicians) through a donor display system (Traxis^SM^) along with relevant secondary donor information such as age, gender, weight, and so on for donor selection. Selection of the best donor among matched alternatives for transplant is based on optimal secondary donor characteristics that provide the best chance of survival for patients. Donor availability remains a key consideration during the donor selection process (Krishnamurti et al. 2003). However, until recently, there have been few metrics to gauge donor commitment and availability. For the purposes of this study, we define donor availability as an adult registry member who responds positively to a Confirmatory Typing (CT) request and successfully completes a blood sample collection.

As a member of a registry only becomes a donor when a stem cell donation is made, and we are interested in predicting the availability of people on the registry, we now use the term member to indicate someone on the registry that we are regarding as a potential donor, and want to investigate their likelihood of being available. Several characteristics are known to be associated with member availability. For example, race and ethnicity are strongly associated characteristics of member availability (G. E. Switzer et al. 2013). There are also established associations with member age, gender, and other characteristics (G. Switzer et al. 1999; Onitilo et al. 2004). Average availability rates are often extrapolated to represent the availability of each member of a given subgroup. Member characteristics positively associated with availability are often not aligned with medical desirability for selection. For example, stem cells from younger male members are known to provide the best chance of survival for patients (Kollman et al. 2016) even though older female members are known to be generally more available. These univariate associations and contradictory relationships make the task of member selection a very difficult and complex process. Choosing a member who will ultimately decline the CT request causes significant delays and complications to completing a transplant, for patients at a time of critical risk for survival. Until recently, clinicians have had to rely on knowledge of historical average availability rates among member subgroups to determine the number of potentially matched members to request for CT to ensure an available member at transplant. A computational model that predicts member availability during the member search process could potentially simplify and improve transplant physicians’ decision process and minimize the risk of delays in transplantation.

In this study, we show how a machine learning approach can be leveraged to predict member availability based on demographics and non-genetic member related factors. Machine learning involves a *learning model* based on historical data to make predictions on future data based on similar patterns. We use *supervised learning* techniques to train models by providing relevant member information as input and their corresponding responses to CT requests as target outputs. The learning techniques learn a mathematical function based on the historical input-output (member information-CT request response) relationship that can be used to predict responses of future CT requests for every member in the registry.

## 2. Data and Methods

Analysis of available data over time has helped experts identify subgroups with higher (or lower) than average availability rates. However, we do not currently have a computational model that unifies all member information (associated with availability) to predict availability. Apart from the demographic data collected at the time a member joins the registry, the NMDP captures several specific member responses and actions associated with member availability. These include:

a. Response to questions on a Post Recruitment Survey that has been specifically developed through research to measure member commitment to the donation process
b. Member responses collected from email and social media invitations to renew commitment
c. Answering a Health History Questionnaire when the member has been identified as a potential match on daily generated search reports and contacted by NMDP personnel
d. Member initiated contact with the NMDP to request updates to their contact information
e. Joining the registry through online registration (versus live drive recruitment where outside influences can more easily sway a person’s decision to join the registry).

It has also been observed that race and ethnicity is a strong indicator of availability. The standard procedure for analyzing the effect of these characteristics on availability is to calculate the historical average among the members belonging to a sub-population (for e.g., members who are self-identified European, or members who have renewed commitment) and extend this average to every member in the population. These averages are also used in studies that assess match rates for different patient populations. Average availability rates for African-Americans have historically been much lower than for Europeans. Gragert et al. (2014) reported average availability rate for European registry members to be 51% and for African-Americans it was only 23% when the study was performed.

We evaluated CT request data from the period August 1, 2013 to November 30, 2015. The responses from all outreach programs had been collected during this period. A total of 178,249 CT requests were made during this period. Associated member data (demographics and response to outreach programs) were collected for modeling. A description of the data collected is listed below.

1. *Member Race and Ethnicity:* every member is assigned to one of the following categories based on self-identified race and ethnicity at the time of registration. To aggregate data over the history of recruitment questionnaires we have assigned members to the following broad race/ethnicity categories. The following race groups are retained as levels in a categorical variable for modeling:

AFA: African-American
API: Asian and Pacific Islander
DEC: Declined to Answer
EUR: European
HIS: Hispanic
MLT: Multi-Race groups
NAM: Native American
OTH: Others
UNK: Unknown
2. *Member Age at request:* Calculated from member birth date and the date the CT request was placed. This information is used as a numeric variable for modeling.
3. *Years on Registry:* Calculated from member registration date and the date the CT request was placed. This information is also used as a numeric variable.
4. *Member Gender:* Collected at the time members join the registry. A binary variable is formed to indicate Female or Male members.
5. *Recommitted:* Registry members are sent communications asking them to renew their commitment to the cause of donation. Whether a member responds to outreach, or not, is recorded in the NMDP database. A response to at least one request is associated with higher availability. A binary response variable is generated for modeling indicating a positive value if they were invited to recommit and responded positively, otherwise negative if they were not invited or they were invited but did not respond.
6. *Address Change:* Identifies member-initiated address change requests. A binary indicator variable is used to identify members who have ever initiated a change of address without solicitation.
7. *Do-It-Yourself (DIY):* Primary method of member recruitment is via live drives where a representative (or a volunteer) from Be The Match^®^ collects information and adds people to the registry. Another way to register is via an online registration, which allows members to register themselves. We have observed that members that register online have a higher availability rate. A binary variable indicates if a member chose to join the registry via the online registration form.
8. *Health History Questionnaire (HHQ) response:* Indicates that the member appeared as a match on a URD search report, was contacted prior to a CT request and answered the health history questionnaire. A HHQ response is also indicative of higher availability. A binary indicator was used for modeling.
9. *Post Recruitment Survey (PRS):* Once a member is added to the registry, a survey is sent to evaluate their commitment to the registry. Members answer four questions about the stem cell donation process that have been found, through research (unpublished), to be most relevant to availability. Members are scored according to their responses. Responses from each of the questions were changed to a positive or negative response. A total composite score was also calculated from answers to the four questions and converted to a positive or negative response. An associated binary variable indicates if a registered member was asked to respond to this survey or not. Another indicator variable was used to identify members who responded to the survey. Each of these variables (2 indicators and 4 response components and an overall response score) were used for modeling. See Appendix A for additional details on the survey.
10. *Donor Center (DC):* The NMDP database contains members recruited through multiple donor center networks. In all there were 13 donor centers used for modeling. These were further grouped into 5 donor center networks. See Table 6 in Appendix B for a list of DC and their network associations.
11. *CT Request Response:* Member responses to CT requests were indicated as either yes or no (declined). The CT request response can be categorized as a “no” request for a number of reasons – unable to contact, medically unavailable, or not interested. This response is used as the *target* variable.

Each of these identified member demographics and responses are associated with varying levels of availability. Figure 1 shows observed availability rates by race for the analysis period covered. Average availability rates for all of the above listed factors are tabulated in Appendix B. Most member features are displayed on the search reporting system (Traxis^SM^) so that transplant centers can select the best member among HLA matched alternatives. This display might indicate some measure of bias in member selection, but the selection process does not directly affect the member availability. We also include certain features in the model that are not displayed but are known to affect availability.

**Figure 1:**
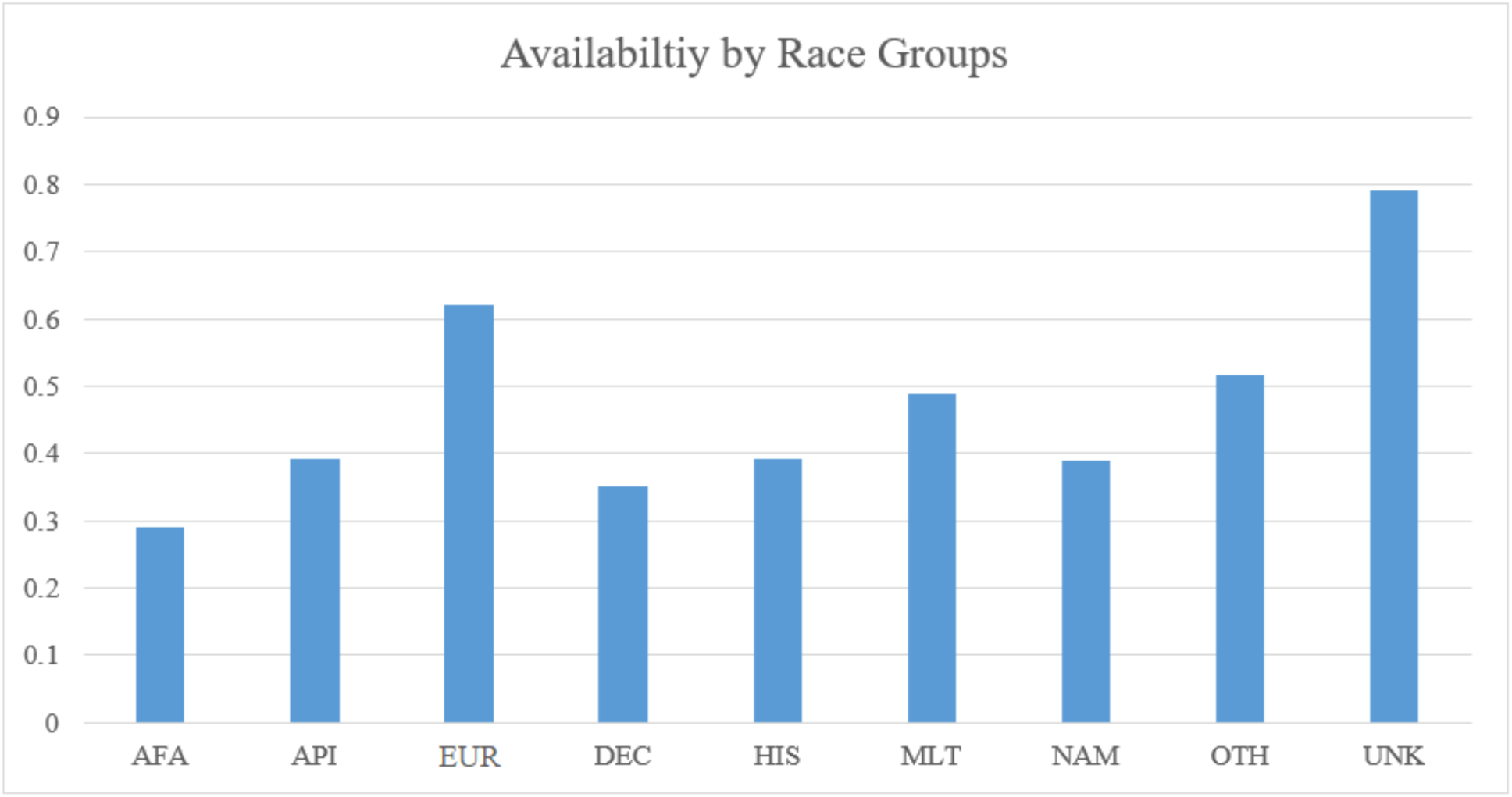
Availability by self-identified broad race groups. Substantial variation is apparent in the average availability rates between groups. Europeans (EUR) have an availability rate of 62% while African-American members have an availability rate of 29%.

Race is often considered a difficult factor to account for in matching, due to errors in self-identified information and complex ancestry information (Hollenbach et al. 2015). Table 1 shows variations in responses to outreach programs by race groups and gender. We notice member responses to outreach programs are not uniformly distributed across either gender or race groups. Traditional determination of availability is done by sub-setting data, for example, into groups such as *EUR Males* from *DC Code A* who have a *Recommit Response* and subsequently calculating the group averages. The number of such possible sub-groups grows swiftly with the number of variables considered. For example, there are 468 possible interactions between Race, DC Code, DIY and HHQ response variables. Sub-setting data into finer detail might show spurious results. We have thus avoided presenting marginal availability values at this level. We note in Appendix B that different variables have different levels of associated average availability. There is also a problem of missing information for members who are sourced from external donor centers. For example, most international members have Race listed as Unknown. When experts have to rely on group averages for availability averages, missing or unreliable information distorts group estimates. Machine learning methods have mechanisms to account for these interactions without users having to specifically define these interactions between variables.

**Table 1:**
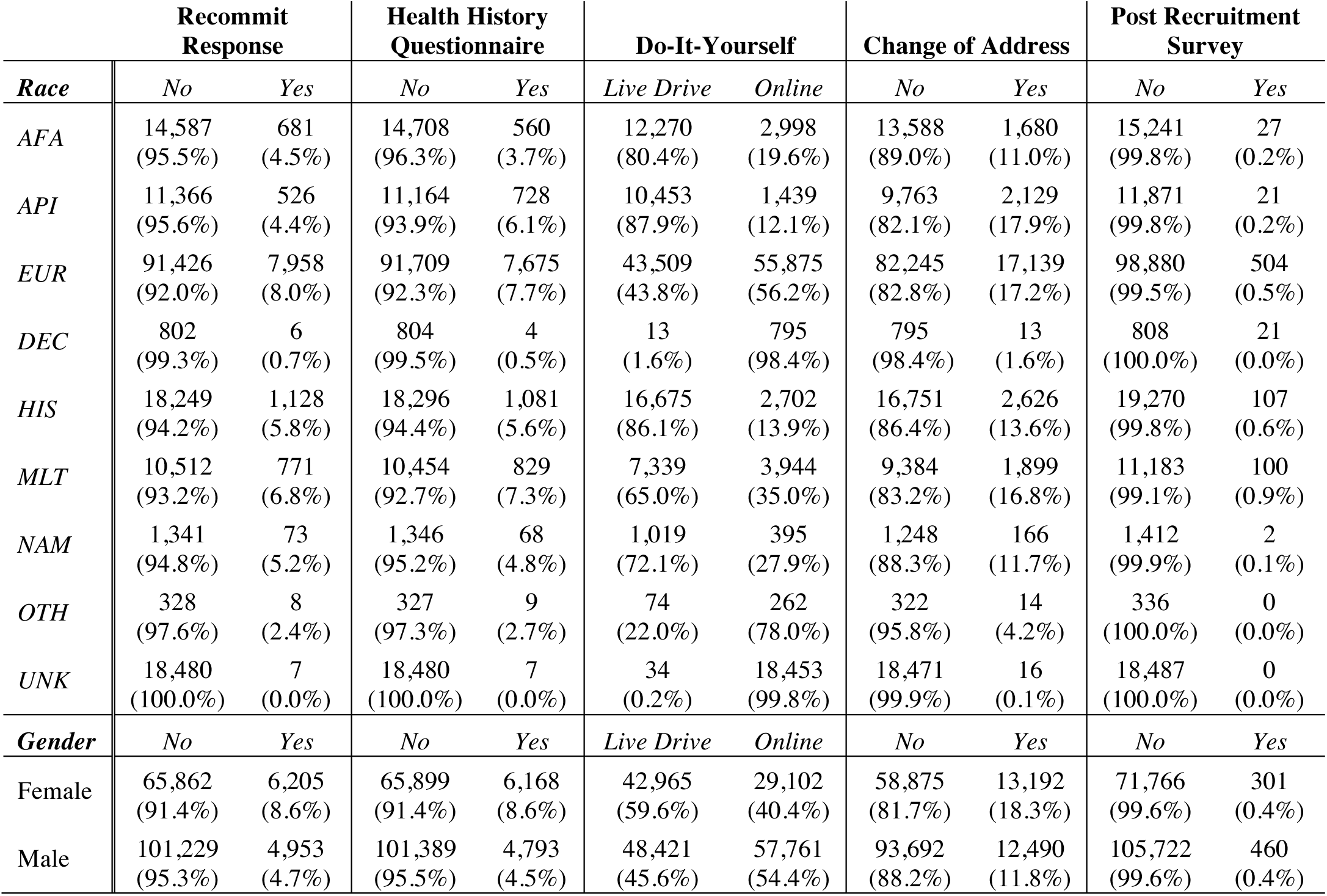
Number of registry members responding to outreach programs and action items broken down by self-identified Race groups and Gender.

|  | Recommit Response |  | Health History Questionnaire |  | Do-It-Yourself |  | Change of Address |  | Post Recruitment Survey |  |
| --- | --- | --- | --- | --- | --- | --- | --- | --- | --- | --- |
| *Race* | *No* | *Yes* | *No* | *Yes* | *Live Drive* | *Online* | *No* | *Yes* | *No* | *Yes* |
| *AFA* | 14,587<br>(95.5%) | 681<br>(4.5%) | 14,708<br>(96.3%) | 560<br>(3.7%) | 12,270<br>(80.4%) | 2,998<br>(19.6%) | 13,588<br>(89.0%) | 1,680<br>(11.0%) | 15,241<br>(99.8%) | 27<br>(0.2%) |
| *API* | 11,366<br>(95.6%) | 526<br>(4.4%) | 11,164<br>(93.9%) | 728<br>(6.1%) | 10,453<br>(87.9%) | 1,439<br>(12.1%) | 9,763<br>(82.1%) | 2,129<br>(17.9%) | 11,871<br>(99.8%) | 21<br>(0.2%) |
| *EUR* | 91,426<br>(92.0%) | 7,958<br>(8.0%) | 91,709<br>(92.3%) | 7,675<br>(7.7%) | 43,509<br>(43.8%) | 55,875<br>(56.2%) | 82,245<br>(82.8%) | 17,139<br>(17.2%) | 98,880<br>(99.5%) | 504<br>(0.5%) |
| *DEC* | 802<br>(99.3%) | 6<br>(0.7%) | 804<br>(99.5%) | 4<br>(0.5%) | 13<br>(1.6%) | 795<br>(98.4%) | 795<br>(98.4%) | 13<br>(1.6%) | 808<br>(100.0%) | 21<br>(0.0%) |
| *HIS* | 18,249<br>(94.2%) | 1,128<br>(5.8%) | 18,296<br>(94.4%) | 1,081<br>(5.6%) | 16,675<br>(86.1%) | 2,702<br>(13.9%) | 16,751<br>(86.4%) | 2,626<br>(13.6%) | 19,270<br>(99.8%) | 107<br>(0.6%) |
| *MLT* | 10,512<br>(93.2%) | 771<br>(6.8%) | 10,454<br>(92.7%) | 829<br>(7.3%) | 7,339<br>(65.0%) | 3,944<br>(35.0%) | 9,384<br>(83.2%) | 1,899<br>(16.8%) | 11,183<br>(99.1%) | 100<br>(0.9%) |
| *NAM* | 1,341<br>(94.8%) | 73<br>(5.2%) | 1,346<br>(95.2%) | 68<br>(4.8%) | 1,019<br>(72.1%) | 395<br>(27.9%) | 1,248<br>(88.3%) | 166<br>(11.7%) | 1,412<br>(99.9%) | 2<br>(0.1%) |
| *OTH* | 328<br>(97.6%) | 8<br>(2.4%) | 327<br>(97.3%) | 9<br>(2.7%) | 74<br>(22.0%) | 262<br>(78.0%) | 322<br>(95.8%) | 14<br>(4.2%) | 336<br>(100.0%) | 0<br>(0.0%) |
| *UNK* | 18,480<br>(100.0%) | 7<br>(0.0%) | 18,480<br>(100.0%) | 7<br>(0.0%) | 34<br>(0.2%) | 18,453<br>(99.8%) | 18,471<br>(99.9%) | 16<br>(0.1%) | 18,487<br>(100.0%) | 0<br>(0.0%) |
| *Gender* | *No* | *Yes* | *No* | *Yes* | *Live Drive* | *Online* | *No* | *Yes* | *No* | *Yes* |
| Female | 65,862<br>(91.4%) | 6,205<br>(8.6%) | 65,899<br>(91.4%) | 6,168<br>(8.6%) | 42,965<br>(59.6%) | 29,102<br>(40.4%) | 58,875<br>(81.7%) | 13,192<br>(18.3%) | 71,766<br>(99.6%) | 301<br>(0.4%) |
| Male | 101,229<br>(95.3%) | 4,953<br>(4.7%) | 101,389<br>(95.5%) | 4,793<br>(4.5%) | 48,421<br>(45.6%) | 57,761<br>(54.4%) | 93,692<br>(88.2%) | 12,490<br>(11.8%) | 105,722<br>(99.6%) | 460<br>(0.4%) |

### Problem Formalization

An important step in building machine learning models is *Problem Formalization*. Here, based on the available data, the modeling task is represented as a learning problem. Our goal is to train a machine learning model to predict member availability. We chose to frame the problem as one of *binary classification* into two groups:

1. *Positive response to a CT request (Available)*
2. *Negative response to a CT request (Not Available)*

To develop a patient-focused model we are considering overall availability, not sub-categories of negative responses such as medical deferral or inability to contact.

In the collected dataset, the overall availability rate is 56%, meaning there was a positive response to 56% of the CT requests made between 8/1/2013 and 11/30/2015. In all, we have 16 variables to represent each member, which includes several categorical variables with multiple levels. Several Machine Learning algorithms were used for modeling - Gradient Boosted Trees (GBM) (Freund and Schapire 1997), Support Vector Machines (SVM) (Cortes and Vapnik 1995) and Logistic Regression. All three techniques are popular learning methods that are commonly used for modeling binary classification formalizations. Logistic Regression works by learning the probability of binary target variable. SVM is a non-probabilistic classification method that learns a decision boundary that separates the two classes (for binary classification). Both SVM and Logistic Regression have been popularly used in multiple biomedical applications including cancer detection and transplant survival analysis studies (Chhatwal et al. 2009; X.-J. Fan et al. 2012; Verplancke et al. 2008; Shiao and Cherkassky 2013; Nematollahi et al. 2017). Gradient Boosting, which falls under a general class of methods called Boosting methods (Breiman 1997; Friedman 2001; Schapire et al. 1998; Freund and Schapire 1997), also learn a decision function. However, unlike the previous two methods, Boosting methods learn multiple classifiers (Decision Trees in this case) on different versions of the training data and eventually combine them to form a single decision rule that has better accuracy than the individual components. Different versions of Boosting techniques have been successfully used in several biomedical studies (Xu et al. 2017; Veta et al. 2015; Ochs et al. 2007). Classification trees (base components in boosting) can handle mixed data (numeric and categorical) and it has practical benefits for our problem.

Modeling was done in R using the *gbm* package (Ridgeway 2006) for Gradient Boosting and the *e1071* (Chang and Lin 2013) and *LiblineaR* (R.-E. Fan et al. 2008) for SVM and Logistic Regression.

## 3. Results and Discussion

The effectiveness of the learning techniques is measured by classification accuracy on an independent test dataset (not used for training the model). The entire dataset is randomly split into two parts: training set and testing set. The training set is used for selecting optimal model parameters using cross validation and the test set is used only to measure performance (accuracy) of the optimal model. Classification Accuracy is evaluated by the percentage of data samples classified correctly by the algorithm. Modeling accuracy results measured on the training and test data are tabulated in Table 2 for the algorithms used for modeling. Details of the training process and model selection criteria are described in Appendix C.

**Table 2:** Training and Testing Accuracy of models measured on the availability dataset.

|  | *Training Accuracy* | *Testing Accuracy* |
| --- | --- | --- |
| *Logistic Regression* | 0.64 | 0.62 |
| *Linear SVM* | 0.65 | 0.63 |
| *Boosted Trees* | 0.73 | 0.70 |

Binary classifiers are commonly used to assign hard labels *(available* or *not available*) on data instances. For this application, it is more beneficial to assign a real-value that can be used to identify availability. Boosted tree methods provide a normalized score for each member in the range of 0–1. From Table 2, we notice it has better classification accuracies than the other two. Hence, we use this method for further analysis.

The *gbm* package provides a variable importance plot from the model that is estimated by calculating the classification improvement provided by each variable split in the trees in the model. A detailed theory about this calculation is provided in Hastie, Tibshirani, and Friedman (2001). Figure 2 shows the relative influence of variables in predicting member availability in the trained model. Only nine of the most important features are shown in the figure.

**Figure 2:**
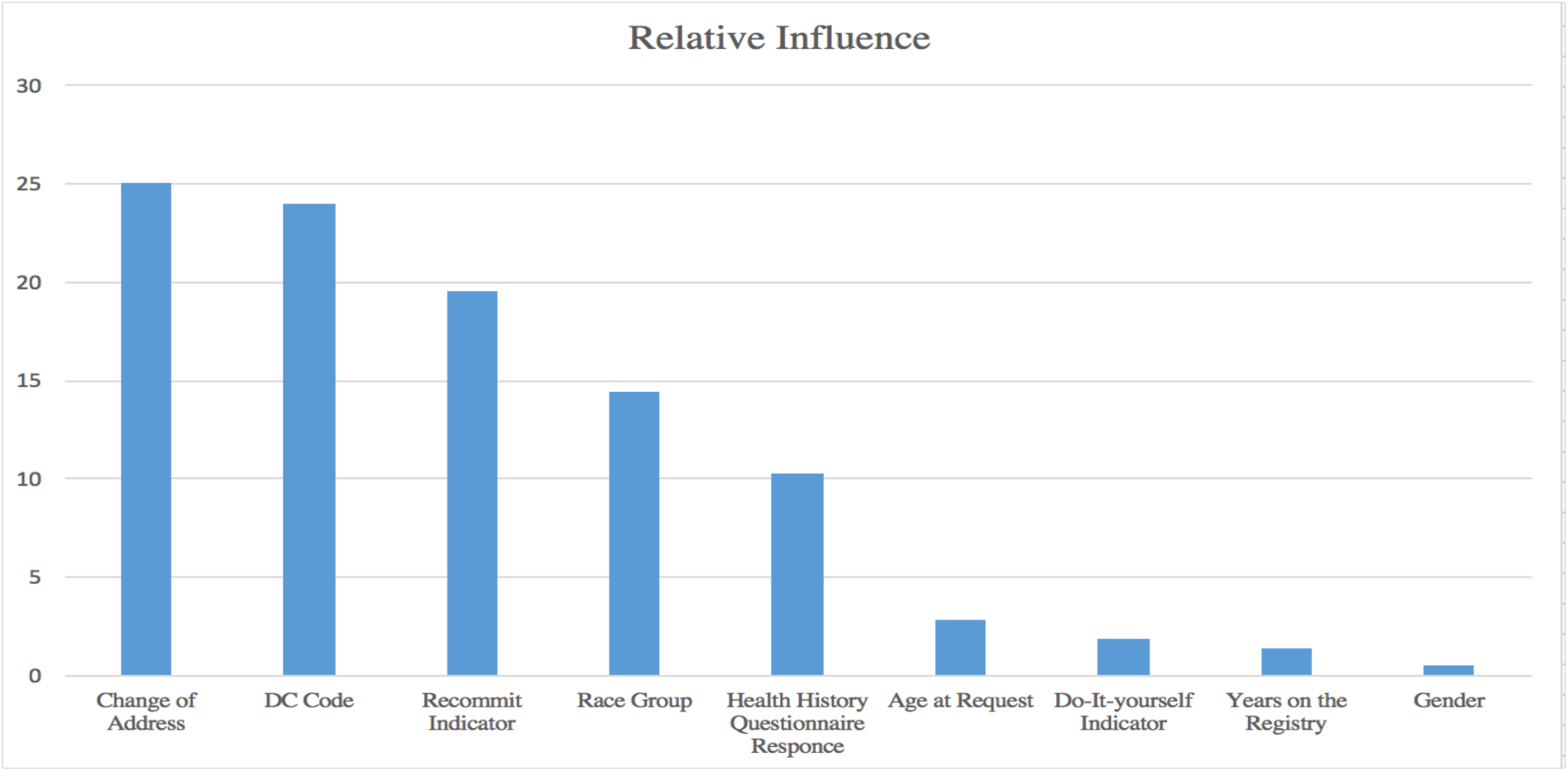
Variable Importance plot for the trained model. Other variables listed in Section 2 had lower influence and are not plotted here.

### Comparing actual Availability to model outcomes

Figure 3 shows actual availability rates broken down by model assigned score ranges. The gradient boosting model assigns a number between 0 and 1 to every member. We have binned members based on these assigned scores and measured the actual availability rate within each bin. For example, availability among members who had a score between 0.9 and 1.0 was 93%. Availability rates in each score range are marked by the rhombus shaped points in the figure. This linear relation between average availability and model assigned scores can be observed across all brackets, as shown in Figure 3. This indicates the model assigned scores linearly correspond to the observed availability. Consequently, model assigned scores can be used as a direct indicator of a member’s availability. The bar for each score range represents the number of members in that range. Average availability and number of members in each score range are represented on different scales on the left and right side of the figure respectively. The linear trend between model scores and actual availability suggests that the model assigned scores are very strong indicators of availability.

**Figure 3:**
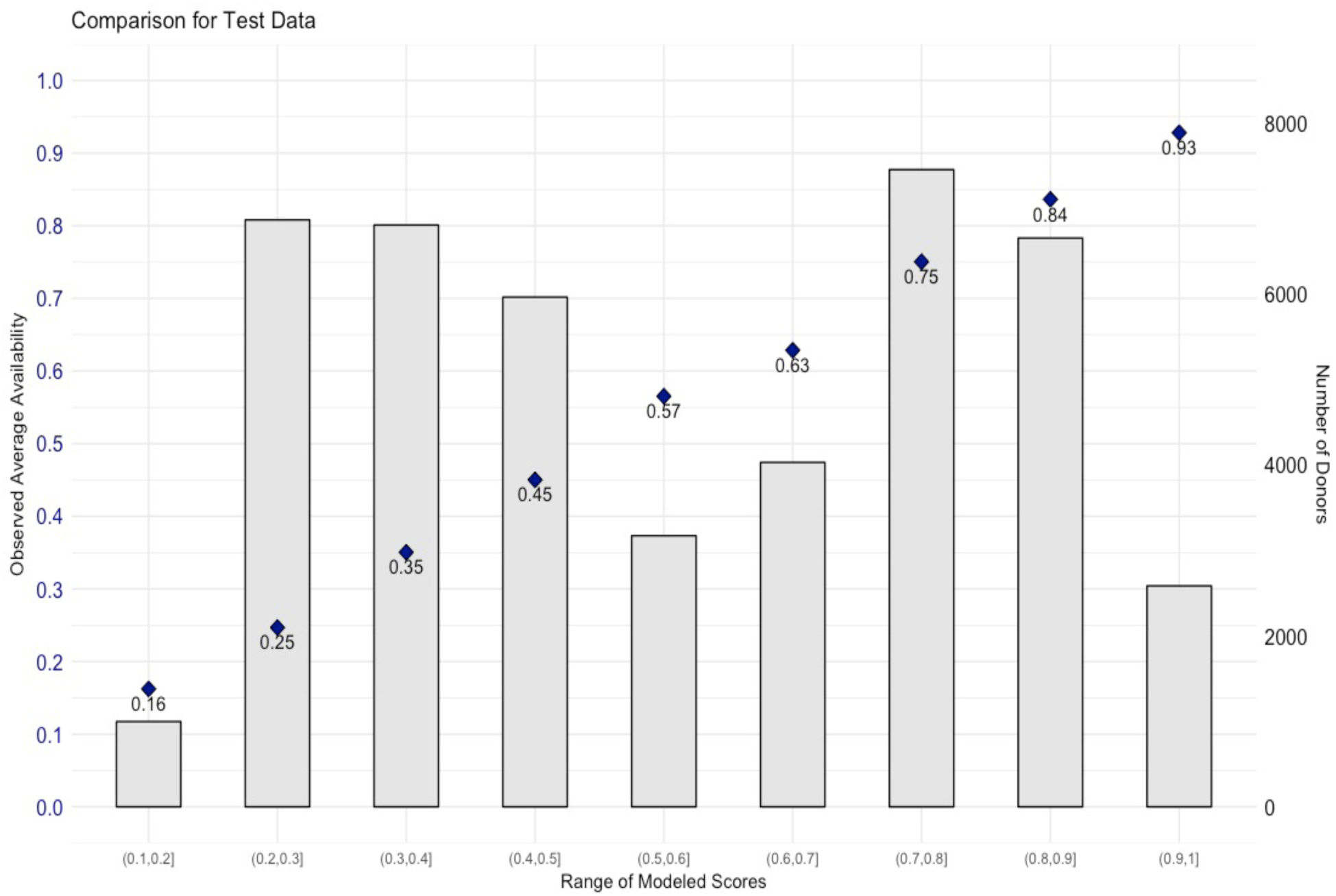
Observed availability rates compared to model assigned scores. The overlaid number in each column represents the average observed availability of members with modeled scores in the corresponding range. The bar graphs are the number of members who are in the brackets noted on the x-axis.

**Table 3.** Example of two registry members input variables and predicted availability.

| Model Variable | Member 1 | Member 2 |
| --- | --- | --- |
| Donor Center Category | 1 | 1 |
| Years on Registry | 3 | 3 |
| Age | 44 | 21 |
| Gender | M | F |
| Race | HIS | AFA |
| Online Registration | Y | N |
| Health History Questionnaire | Y | N |
| Recommitted | Y | N |
| Address Change | 0 | 0 |
| Post-recruitment Survey Sent | 0 | 0 |
| Post-recruitment Survey Reply | NA | NA |
| Post-recruitment Survey Score | NA | NA |
| Predicted Availability | 0.95733 | 0.13387 |

The NMDP provides access to the US Be The Match Registry as well as several other domestic and international registries. Responses to outreach programs are only recorded for the Be The Match recruits. This results in different levels of information available for modeling availability across donor center networks. The model is capable of adjusting for varied levels of information and assigning a score to every member in the registry.

Figure 4 shows the density of availability scores across each of the five DC networks. Since we have access to more model inputs for the registry members in Be The Match (Network 1) the model is able to provide greater differentiation. For other networks, the lack of inputs results narrower distribution of values.

**Figure 4:**
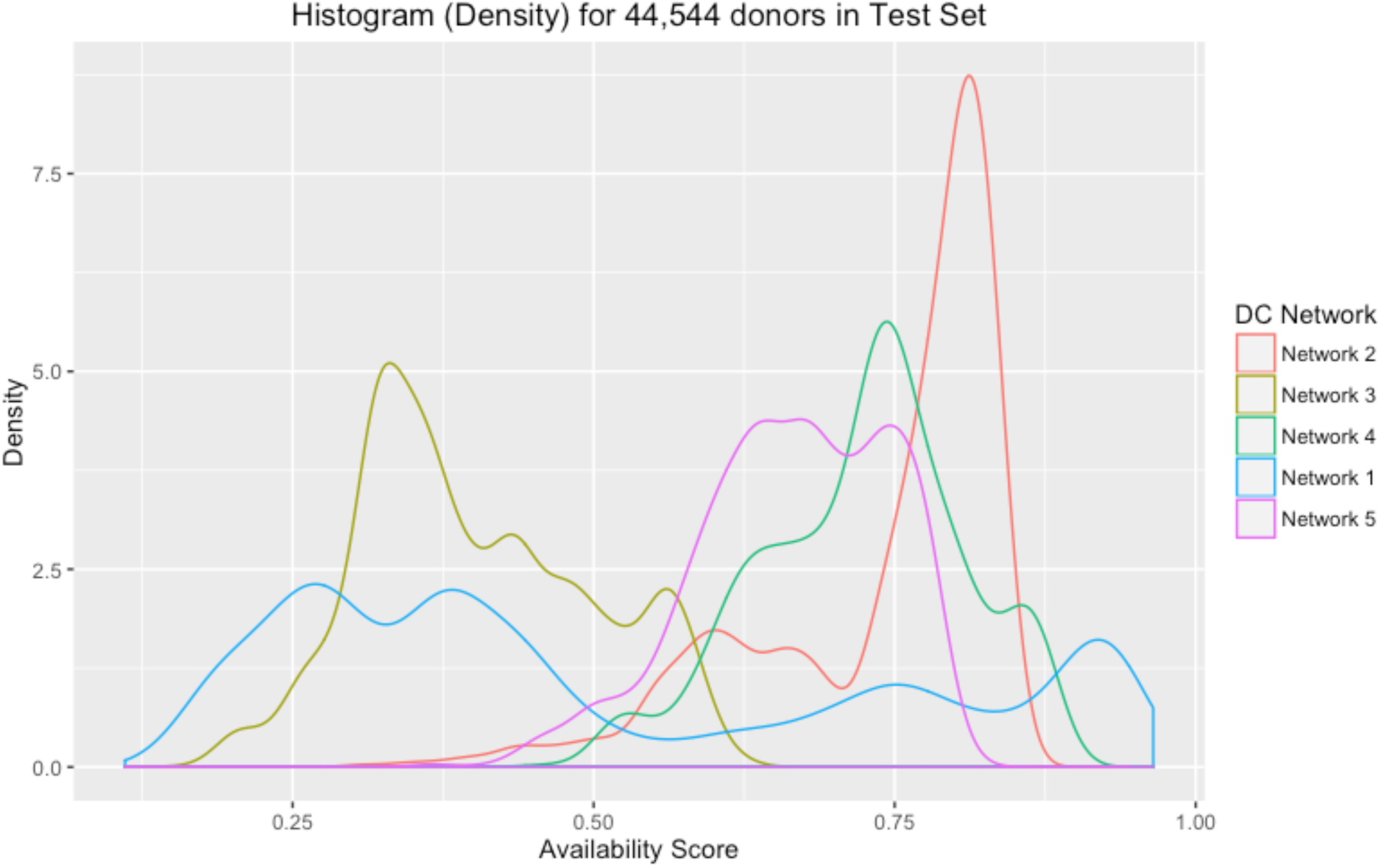
Density of member availability for different donor networks

## 4. Conclusion

Typically, experts have to rely on historical averages for availability estimates. This results in inaccuracy in estimating an individual member’s availability. This also limits the ability of clinicians to accurately estimate how many members should be contacted in order to ensure the patient will be supplied with a bone marrow transplant product when needed. The proposed system helps by providing an individual estimate for each member based on the set of factors available for estimating their availability. Creating a single score for each potential member can significantly simplify the member selection process and reduce the time taken to successfully complete the transplant.

## Acknowledgement

The methods used for this analysis were developed through a research grant funded by the US Office of Naval Research (N00014-17-1-2388). We thank Bronwen Shaw, Juliet Barker, Stephen Spellman, Jason Dehn, and Michael Wright for their valuable comments and suggestions on this manuscript.

## Appendix A

### Post Recruitment Survey Inputs

#### The four post recruitment survey questions

1. Imagine that we are calling you today to ask you to give a blood sample for further testing to see if you are the best match for a patient. On a scale of 1 to 5 where 1 is very likely and 5 is not at all likely, how likely are you to agree to give a blood sample?
2. If you are selected as a match for a patient, the total time commitment for testing and actual donation is about 40 hours over a 4–6 week period. On a scale of 1 to 5, how willing would you be to make that time commitment to donate?
3. If further testing showed that you were the best match for a patient, on a scale of 1 to 5, how likely is it that you would actually donate?
4. On a scale of 1 to 5 where 1 is very willing and 5 is not at all willing, how willing would you be to donate to someone you don’t know?

Responses of 7 (don’t known) and 8 (left blank) were also possible

#### Encoding responses to questions

Responses of 1 and 2 were encoded as positive Responses of 3–5 were encoded as negative. Responses of 7 and 8 were encoded as negative

#### Composite Scoring

Calculating the composite score from the four Post Recruitment Survey (PRS) questions:

PRS response is positive if the composite score is 4. PRS response is negative if the composite score >4 (maximum score would be 8).

If response to a question is 1 or 2, it is counted in the composite calculation below as a 1. If response to a question is 3, 4 or 5, it is counted in the composite calculation below as a 2.

If two or more responses are missing then the PRS response is negative. If there is only one response missing, then that response is a 1 in the composite calculation below. A response of 7 or 8 is the same as a missing score.

PRS composite score = (Converted as indicated above) Sum of converted responses to questions 1 through 4.

## Appendix B

Marginal Availability rates for different factors listed in Section 2 are tabulated below. Marginal availability rates are affected by sample size for certain factors. We include all of these in the modeling for their assumed effect on availability.

**Table 4:** Marginal Availability rate by Gender.

| ***Gender*** | ***Availability Rate*** | ***Count*** |
| --- | --- | --- |
| Male | 55.8% | 106,182 |
| Female | 55.9% | 72,067 |

**Table 5:** Marginal Availability by Health History Questionnaire Response.

| ***Questionnaire Response*** | ***Availability Rate*** | ***Count*** |
| --- | --- | --- |
| Responded | 87.1% | 10,961 |
| Not Responded | 53.8% | 167,288 |

**Table 6:** Marginal Availability Rates by Donor Centers.

| ***Donor Center Code*** | ***Network Name*** | ***Network Code*** | ***Availability Rate*** | ***Count*** |
| --- | --- | --- | --- | --- |
| A | NMDP | 1 | 50.4% | 100,057 |
| B | INT | 4 | 67.6% | 470 |
| C | INT | 4 | 62.1% | 863 |
| D | DOD | 3 | 39.9% | 22,140 |
| E | DKMS | 2 | 79.6% | 31,332 |
| F | INT | 4 | 69.3% | 990 |
| G | INT | 4 | 72.6% | 212 |
| H | INT | 4 | 76.5% | 243 |
| I | US - NN | 5 | 64.8% | 3,062 |
| J | DKMS | 2 | 58.9% | 13,708 |
| K | US - NN | 5 | 48.8% | 1,695 |
| L | DKMS | 2 | 75.5% | 3,152 |
| M | INT | 4 | 68.7% | 323 |

**Table 7:** Marginal Availability Rate by Post Recruitment Survey Response.

| ***Post Recruitment Survey*** | ***Availability Rate*** | ***Count*** |
| --- | --- | --- |
| Response | 75.2% | 761 |
| No Response | 55.8% | 177,488 |

**Table 8:** Marginal Availability by Recommit Response.

| ***Recommit Request*** | ***Availability Rate*** | ***Count*** |
| --- | --- | --- |
| Response | 93.5% | 11,158 |
| No Response | 53.4% | 167,091 |

**Table 9:** Marginal Availability Rates for DIY record.

| ***Do-It-yourself Registration*** | ***Availability Rate*** | ***Count*** |
| --- | --- | --- |
| Online | 64.6% | 86,863 |
| Others (Donor Drive) | 47.6% | 91,386 |

**Table 10:**
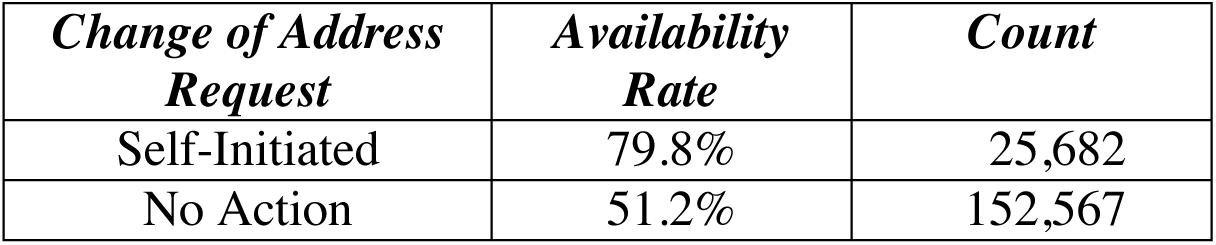
Marginal Availability Rate for Change of Address requests.

## Appendix C

We use binary classifiers (as we have two categories: *available* and *not available*) to model the availability. Classifiers estimate a function *f(****x***) that separates the input space into two regions corresponding to the two classes, where **x** is a feature vector that is used to represent measurements/observations for a data instance. For our problem, **x** is a vector representing member characteristics listed in Section 2. Each member is represented by a vector (row in the dataset). The estimated function is then used to assign labels based on the feature vector. The quality of a classifier is measured by a loss function. If a classifier predicts all the given data instances perfectly the loss will be 0. The loss function measures the discrepancy between predictions by the classifier and the actual desired output. A standard loss function used is the 0/1 function shown in Equation 1.

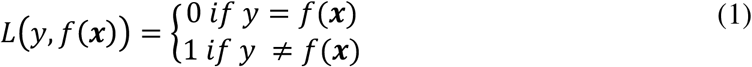

*y* is the actual observed output, *f(****x***) is the model predicted output for sample **x**.

Classifiers typically have tuning parameters that need to be adjusted by the user according to the data being modeled; this is achieved by cross validation. The entire dataset is typically randomly separated into two groups, one for training the model and the other for testing the accuracy of the model. Depending on the size of the dataset available, the split ratio is determined. For our problem, we use a 75:25 split (training: testing). A k-fold cross validation is then performed on the training set to tune model parameters. A grid of parameters is searched, and the best parameter is chosen based on lowest average validation across all folds. We use a 5-fold cross validation to select the best parameter. Depending on the modeling method, users need to tune one or two parameters. After the best parameter is determined, a model is trained on the entire training data split. Classifier accuracy is then measured on the same set and is recorded as the *Training Accuracy/ Error*. However, the quality of prediction is determined by the classifier performance on the test set. None of the samples in the testing set is used for model selection; the test set is only used for assessing model accuracy, *Testing Accuracy/Error*. Error is measured as the average error across all samples in the dataset, as show in Equation 2.

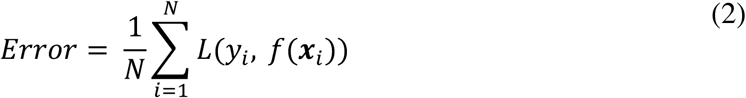

This is the average error for a dataset with *N* samples and the loss function is as defined in Equation 1. While the above-mentioned metric is the most commonly used, care should be taken to use an appropriate metric that addresses specific requirements of the modeling problem. We refer readers to (Cherkassky 2013; Cherkassky and Mulier 2007) for a detailed discussion on model selection procedure and further details on modeling techniques.

### Data Pre-Processing

Raw data is processed before model training and is an important part of the modeling process. Apart from following standard procedures for pre-processing, it is also important to reflect domain knowledge in the pre-processing.

Age at request and Years on registry are scaled on a 0–1 range. The maximum and minimum are recorded to process future samples to the same scale. Categorical predictors (Race and Donor Center Code) with multiple levels are used as *factor* variables in R software. Other predictors, as listed in Appendix B with only two possible values, are encoded as binary variables. Care is taken to identify missing values and miscellaneous responses to outreach requests.

